# Measuring Selection Across HIV Gag: Combining Physico-Chemistry and Population Genetics

**DOI:** 10.1101/204297

**Authors:** Elizabeth Johnson, Michael A. Gilchrist

**Affiliations:** Microbiology, University of Tennessee, Knoxville, TN, United States; National Institute for Mathematical and Biological Synthesis, University of Tennessee, Knoxville, TN, United States; Ecology and Evolutionary Biology, University of Tennessee, Knoxville, TN, United States

## Abstract

We present physico-chemical based model grounded in population genetics. Our model predicts the stationary probability of observing an amino acid residue at a given site. Its predictions are based on the physico-chemical properties of the inferred optimal residue at that site and the sensitivity of the protein’s functionality to deviation from the physico-chemical optimum at that site. We contextualize our physico-chemical model by comparing our model fit and parameters it to the more general, but less biologically meaningful entropy based metric: site sensitivity or 1/*E*. We show mathematically that our physico-chemical model is a more restricted form of the entropy model and how 1/*E* is proportional to the log-likelihood of a parameter-wise ‘saturated’ model. Next, we fit both our physico-chemical and entropy models to sequences for subtype C’s Gag poly-protein in the LANL HIV database. Comparing our model’s site sensitivity parameters *G*′ to 1/*E* we find they are highly correlated. We also compare the ability of *G*′, 1/*E*, and other indirect measures of HIV fitness to empirical *in vitro* and *in vivo* measures. We find *G*′ does a slightly better job predicting empirical fitness measures of *in vivo* viral escape time and *in vitro* spreading rates. While our predictive gain is modest, our model can be modified to test more complex or alternative biological hypotheses. More generally, because of its explicit biological formulation, our model can be easily extended to test for stabilizing vs. diversifying selection. We conjecture that our model could also be extended include epistasis in a more realistic manner than Ising models, while requiring many fewer parameters than Potts models.

## Introduction

HIV’s protein sequence has been described as having a great deal of ‘genetic plasticity’ due to the fact that amino acid substitutions at many different sites appear to have little or no consistent effect on HIV fitness (Lemey et al., 2006; Salemi, 2013; Cuevas et al., 2015; Rihn et al., 2013). Such genetic plasticity allows HIV populations to evade or ‘escape’ the patient’s immune response by simply evolving to a new location in epitope space. HIV’s remarkable ability to escape the host’s immune response has impeded efforts to create an effective vaccine (Goulder and Watkins, 2004; Autran et al., 2008; Johnston and Fauci, 2008).

In response, researchers have tried to identify amino acid sites under strong and consistent stabilizing selection (Ferguson et al., 2013; Liu et al., 2013; Barton et al., 2016a), believing them to be promising vaccine targets (Rolland et al., 2013). In order to identify these sites, researchers have attempted to estimate HIV’s fitness landscape (Deforche et al., 2008; Seifert et al., 2015; Kouyos et al., 2012; Hinkley et al., 2011; Shekhar et al., 2013; Ferguson et al., 2013; Mann et al., 2014; Lorenzo-Redondo et al., 2014; Moradigaravand et al., 2014; Barton et al., 2016a). Since first introduced by Wright (1932), fitness landscapes have become a conceptual cornerstone within the field of evolutionary biology (e.g. Lande and Arnold, 1983; Lande, 1985, 1986; Kauffman and Levin, 1987; Charlesworth and Rouhani, 1988; Kauffman, 1993; Niklas, 1994; Gavrilets, 1997; Fontana, 2002; Berg and Lässig, 2003; Gavrilets, 2004; Berg et al., 2004; Wilke and Drummond, 2006; Gilchrist, 2007; Lässig, 2007; Calcott, 2008; Mustonen and Lässig, 2009; Draghi et al., 2010; Gilchrist et al., 2009; Wallace et al., 2013; Gilchrist et al., 2015). In the simplest scenarios where the evolutionary process has either reached stationarity or we have no *a priori* information, the probability of observing a sequence *i* is proportional to its evolutionary fitness *W*_*i*_ raised to the effective population size *N*_*e*_, i.e. 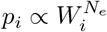 (Wright, 1969; Iwasa, 1988; Berg and Lässig, 2003; Sella and Hirsh, 2005; McCandlish et al., 2015).

While often unaware of its theoretical foundation, HIV researchers have extensively used this link between the fitness contribution of an amino acid residue at a particular site and its observed genotype frequency (e.g. Rihn et al., 2013; Liu et al., 2006). For example, researchers have adapted the Potts and Ising models from statistical mechanics to quantify direct and epistatic fitness effects between amino acid sites (Ferguson et al., 2013; Mann et al., 2014; Barton et al., 2016a). While the Ising model, which categorizes amino acid residues as optimal or non-optimal, has been criticized as being overly simplistic, it has been effectively employed to identify epistatic interactions between sites (Ferguson et al., 2013; Mann et al., 2014). Instead of using a simple optimal/non-optimal categorization, the Potts model categorizes each of the 20 amino acid separately. While this greater categorization makes the Potts model more biologically realistic, it also makes it hard to fit. Specifically, in order to describe epistatic effects between any two sites, the Potts model requires the estimation of hundreds of parameters. As a result, confidently fitting this model to multiple sequence alignments is infeasible, even with large data sets (Barton et al., 2016a). Researchers have also used Shannon entropy *E*, specifically the inverse entropy of a site 1/*E* (i.e. its ‘conservation’), as a measure of the strength of consistent stabilizing selection for the presumably optimal consensus amino acid residue (Dietrich and Skipper, 2012; Acevedo et al., 2014). In this approach, sites with little variation in amino acid usage have high 1/*E* values and are, in turn, inferred to be experiencing strong and consistent stabilizing selection. In contrast, sites with substantial variation have low 1/*E* values and are inferred to be either under weak or variable stabilizing selection. As we show later, the conservation of a site 1/*E* is proportional to the expected log-likelihood of observing a randomly chosen amino acid under a saturated, multinomial model parameterized with a given data set.

Given its definition, site conservation 1/*E* can is best viewed as a summary statistic quantifying the ruggosity of HIV’s fitness landscape (Kauffman, 1993) rather than a direct measure of consistent stabilizing selection for an optimal amino acid. Another shortcoming of 1/*E* as a biologically meaningful metric is the fact that it treats all amino acid residues as being equally dissimilar from the one another. As a result, 1/*E* ignores the fact that amino acid residues have differing degrees of physico-chemical dissimilarity. This is undesirable given that the physico-chemical properties of amino acids clearly affect the probability one amino acid will substitute for another (Grantham, 1974; Wilke and Drummond, 2010). As a result, 1/*E* has the potential to miss sites where there is strong, consistent stabilizing selection for a set of physico-chemical properties, but where these properties can be reasonably satisfied by more than one amino acid residue. More generally, the fact that 1/*E* ignores the physico-chemical properties of amino acid residues suggests it is glossing over information embedded within the data.

Despite the fact that site conservation 1/*E* ignores the physico-chemical characteristics and is not actually a measure of consistent stabilizing selection against non-optimal amino acid residues, it is widely used. One contributing factor to 1/*E*’s wide use is the fact that it can be easily calculated from sequence alignment data (c.f. the Potts model). A second contributing factor to 1/*E*’s popularity is its utility. Even though 1/*E* is a coarse meteric, it has proven useful for identifying ‘fragile’ sites, i.e. those with low genetic plasticity.

In an effort to overcome these aforementioned shortcomings, we introduce a model that is more biologically detailed than 1/*E*, but less parameter rich than the Potts model. First, instead of treating amino acids residues as equally dissimilar categories, our model uses a subset set of physico-chemical properties and weighting terms to describe the differences between residues. Second, it explicitly estimates the sensitivity of viral fitness to deviations from the physico-chemical of the optimal amino acid for a given site. While the site sensitivity parameters *G*′ we estimate correlate well with site conservation 1/*E*, *G*′ does a slightly better job predicting HIV genotype fitness from both *in vivo* and *in vitro* studies. Unlike site conservation 1/*E*, our model allows us to test biologically motivated hypotheses. For example, we find that our physico-chemical weightings and the distributions of *G*′ vary between the different protein regions of Gag.

Taken together, our physico-chemical model helps advance researchers ability to extract information from observational data and represents a biologically grounded framework which can be further extended and used to test clearly posed hypotheses. In its current form, our physico-chemical model only considers site independent effects and can be viewed as a constrained version of the more general, parameter saturated, entropy model underlying the calculation of 1/*E*. Given the parallels between evolution and statistical mechanics (Sella and Hirsh, 2005), it should be possible to extend our physico-chemical model to include epistatic effects in a more realistic manner than the Ising models, which ignore physico-chemical properties, but in a substantially more efficient manner, in terms of the number of epistatic parameters and the ease of parameter estimation, than the physico-chemical informed Potts models.

## Methods

We begin by presenting our physico-chemical model and, in the process, clearly define our site sensitivity parameter *G*′. Next, we review how site conservation 1/*E* is defined using Shannon entropy. We then clearly show the link between the physico-chemical model and the entropy model and their corresponding probabilities of observing each amino acid residue at a particular site. Finally, we describe the data and methods we used to parameterize the physico-chemical and entropy models and evaluate their ability to predict empirical measurements of HIV fitness. Definitions of all of our model parameters can be found in Table 1.

**Table 1.**
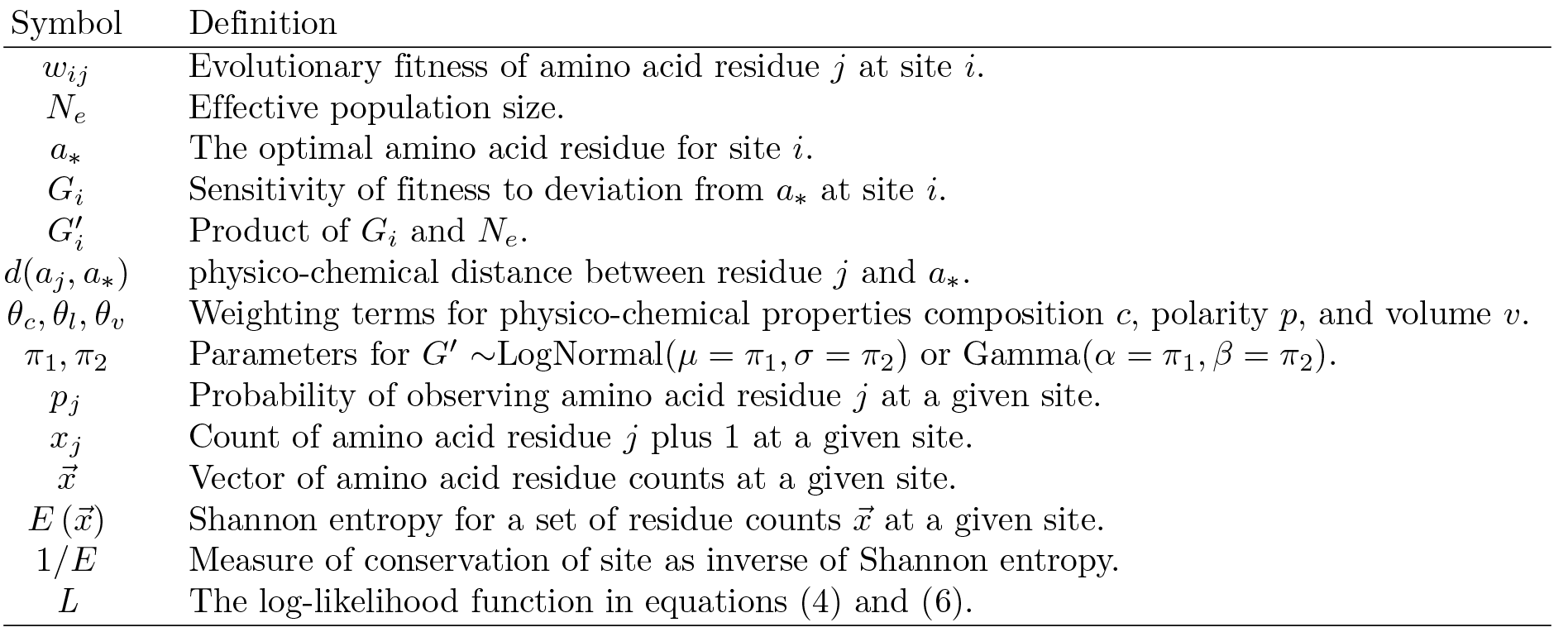
Symbols used in this work and their verbal definitions. Symbol Definition

| Symbol | Definition |
| --- | --- |
| $w_{ij}$ | Evolutionary fitness of amino acid residue $j$ at site $i$. |
| $N_e$ | Effective population size. |
| $a_*$ | The optimal amino acid residue for site $i$. |
| $G_i$ | Sensitivity of fitness to deviation from $a_*$ at site $i$. |
| $G'_i$ | Product of $G_i$ and $N_e$. |
| $d(a_j, a_*)$ | physico-chemical distance between residue $j$ and $a_*$. |
| $\theta_c, \theta_l, \theta_v$ | Weighting terms for physico-chemical properties composition $c$, polarity $p$, and volume $v$. |
| $\pi_1, \pi_2$ | Parameters for $G' \sim \text{LogNormal}(\mu = \pi_1, \sigma = \pi_2)$ or $\text{Gamma}(\alpha = \pi_1, \beta = \pi_2)$. |
| $p_j$ | Probability of observing amino acid residue $j$ at a given site. |
| $x_j$ | Count of amino acid residue $j$ plus 1 at a given site. |
| $\vec{x}$ | Vector of amino acid residue counts at a given site. |
| $E(\vec{x})$ | Shannon entropy for a set of residue counts $\vec{x}$ at a given site. |
| $1/E$ | Measure of conservation of site as inverse of Shannon entropy. |
| $L$ | The log-likelihood function in equations (4) and (6). |

### Modeling the HIV Fitness Landscape

In this study, we focus on two structurally similar models for describing a HIV fitness landscape: our physico-chemical based approach and Shannon entropy. Both models assume the fitness landscape is fixed and that each amino acid site affects viral fitness independent of the others. In the physico-chemical model the expected frequencies of the different amino acid residues are determined by their physico-chemical properties, the optimal residue for that site *a*_*_, and the strength of consistent stabilizing selection *G* for *a*_*_. These expected frequencies also depend on the physico-chemical weights 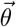, which are shared across a set of sites.

In contrast to our physico-chemical model, in the Shannon entropy model there are no shared parameters between sites, the optimal amino acid residue is assumed to be the most frequent one, and the frequency of the 19 other non-optimal amino acids residues are completely unconstrained and set to equal their observed frequencies for that site. Technically, the Shannon entropy model could be viewed as an unconstrained version of our physico-chemical model. In the Shannon entropy model there are 19 free parameters per site. In contrast, in our physico-chemical model there are only 2, *a*_*_ and *G*, along with a minimum of 3 global parameters describing physico-chemical weights and a hierarchical distribution for *G*. We use AIC for model comparisons.

#### A Physico-Chemical Model

For a given site, we assume the fitness *w* of amino acid residue *a_j_* declines exponentially as a product of a site specific sensitivity parameter *G*_*i*_ and the distance *d*, in physico-chemical space, of *a*_*j*_ from the optimal amino acid residue for that site *a*_*_. That is,

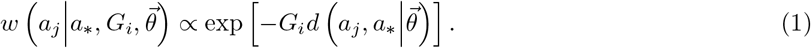

We use the euclidean distance function

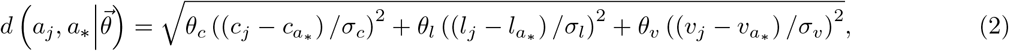

the physico-chemical weighting terms 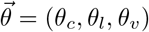 represent *a priori* unknown weights for the known amino acid residue physico-chemical properties: the ratio of carbon to non-carbon atoms or ‘composition’ *c*, polarity *p*, and molecular volume *v*. The physico-chemical weights 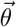 are are assumed to be shared across multiple sites and are estimated from the data. In order to account for the fact that physico-chemical properties are measured in different units and facilitate their interpretation of our weight terms 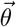, the differences in physico-chemical properties are scaled by the standard deviation of this property observed across the 20 canonical amino acids, i.e. *σ*_*i*_. As a result of this scaling, the weighting term describe the sensitivity of protein function to the deviation in physico-chemical properties between amino acid relative to the total amount of variation possible. Our choice of physico-chemical properties follows Grantham (1974); other physico-chemical properties could be used instead (Sharma et al., 2013). Because *G*′ is always multiplied by the distance function *d* in our physico-chemical model, there is an inherent lack of identifiability of these terms. To solve this problem and facilitate ease of interpretation, 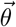 was constrained so that the sum of its values equaled 1.

Following the example of other researchers in this field (Ferguson et al., 2013; Mann et al., 2014; Barton et al., 2016a) we treat HIV sequence data from different patients as independent samples from the evolutionary stationary distribution as described by Sella and Hirsh (2005). We define the composite parameter 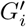 as the product of the site sensitivity and the effective population size *N*_*e*_, i.e. 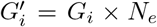. For now we ignore the effects of mutation bias. In order to avoid issues that would result from the infinite MLE of *G*′ at invariant sites, we employed hierarchical approach where *G*′ is assumed to follow either a LogNormal or Gamma distribution. As a result, the parameters *π*_1_ and *π*_2_ represent the shape and scale parameters or the log scale mean and standard deviation for *G*′, respectively. We allowed the *G* distributions and parameters to be shared or vary between each of the gag poly-peptide regions: the nucleo protein p6, nucleocapsid protein p7, matrix protein p17, and capsid protein p24. As a result, the probability *p* of observing amino acid residue *j*, at site *i* is,

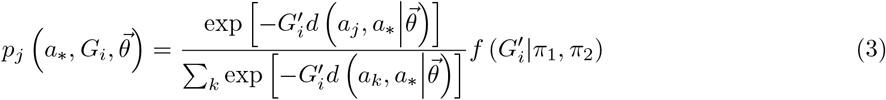

Based on our model’s assumptions and structure, 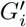 represents a quantitative measure of the strength and efficacy of consistent stabilizing selection on a given site relative to genetic drift.

Given these assumptions, the probability of observing a set of amino acid residue counts 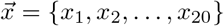 at a given site *i* follows a multinomial distribution. That is,

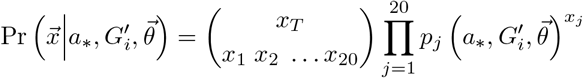

where *x*_*T*_ = Σ_*j*_ *x*_*j*_ is the total number of observations made at site *i* and and *a*_*_ is the optimal amino acid for site *i*. Correspondingly, the Log-Likelihood *L* of the data 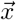 as a function of our physico-chemical the model parameters, *a*_*_, 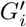, and 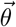 is,

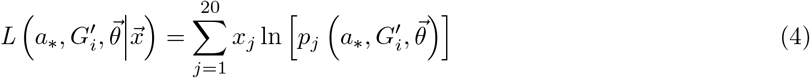

By maximizing Eq.(4) with respect to our model parameters, we can identify the most likely parameter values and our confidence in them: the physico-chemical weights 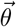, the optimal amino acid residue at each site *a*_*_, and the *N*_*e*_ scaled sensitivity of HIV fitness to deviations in physico-chemical space for each site *G*′. By linking fitness to the physico-chemical qualities of an amino acid residue, we effectively reduce the number of parameters in our multinomial model from 19 parameters per site to 2 parameters per site, 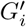 and *a*_*_ plus, depending on how we partition the data, 2 or 8 shared physico-chemical weight parameters 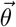. As a result, our physico-chemical model is highly ‘unsaturated’.

#### The Entropy Model

While Shannon entropy is a measure of information of a message, it is also proportional to the log of the probability of the data 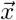 under a multinomial model where the probability of each category *p*_*j*_ is equal to its observed relative frequency *x*_*j*_/*x*_*T*_ where, as before, *x*_*T*_ is the number of observations. Setting *p*_*j*_ = *x*_*j*_/*x*_*T*_ not only makes intuitive sense, it is also equal to the maximum likelihood estimate (MLE) of *p*_*j*_ under a multinomial model. Thus Shannon entropy of a given site *i* with a set of counts 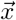 can be written as

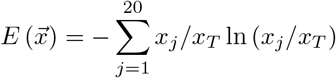

and is related to the probability of the data 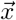 at the maximum likelihood values of its parameters 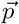 under a multinomial model

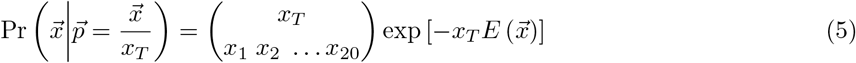

Correspondingly, the maximum Log-Likelihood of the data 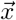 under the Shannon entropy model is,

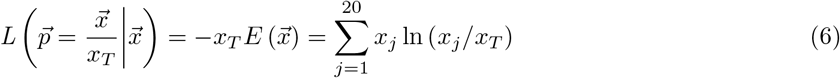

The entropy of a site 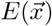 can be interpreted in a number of different ways. One interpretation of 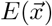 is as a diversity index for amino acid residues at a site or region (Jost, 2006). Another interpretation is, under certain conditions, 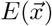 is proportional to the expected number of guesses one must make to infer the state of a site. Equivalently, 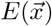 is also equal to the the mean contribution of a category to a site’s log-likelihood.

Despite having many potential interpretations, because of its descriptive and ‘many to one’ nature, there is no clear way of linking the Shannon entropy of a site *E* and the strength of purifying selection for or against a particular amino acid. Finally, the inverse of *E* is used to describe the conservation of a site or region 1/*E* which has, in turn, been used as a heuristic measure of the strength of consistent stabilizing selection for the optimal amino acid residue at a site (Allen et al., 2005; Liu et al., 2012a). In addition, because the sum of *p*_*i*_ must equal one, from a likelihood perspective the entropy model has 20 − 1 = 19 free parameters per site, making it a fully saturated model.

### Data

#### Amino Acid Sequences

To parameterize the models we used the Gag poly-peptide of HIV subtype C MSA. We excluded linker regions and, as a result, analyzed 520 sites using 1058 curated sequences from the Los Alamos National Laboratory (LANL) database (Biophysics Group Los Alamos National Lab, 2016). This database contains filtered web alignments curated by the biophysics group at LANL. Details concerning the curation and filtering of the sequences in the alignment process can be found at http://www.hiv.lanl.gov/. Data was downloaded in September 2016. Sequences were processed to obtain counts for amino acid residue *j* at site *i* for each HXB2 site.

#### Empirical Evaluation of Model Predictions

In order to test whether our site sensitivity parameter *G*′ is more biologically informative than the standard conservation metric 1/*E*, we evaluated its ability to predict empirical observations of *in vivo* and *in vitro* viral fitness. To test our physico-chemical model’s ability to predict *in vivo* behavior of HIV, we used viral escape times from a patient’s CD8 T-cell response as estimated by Barton et al. (2016a) using the patient data from Liu et al. (2012b). In that study, escape times of viral epitopes from the patient’s immune response were estimated using sequential sequencing data from reactive epitopes sites in multiple HIV-1 subtype B and C infected patients over 3 years. These researchers found that that the mean conservation value 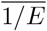 for an epitope is positively correlated with escape time. Epitopes were defined as 8-11 amino acid long regions of the proteome recognized by patient specific T cell immune response. Escape time *t* was defined as the number of days between detection of the T-cell response and the time viral variants bearing that respective reactive epitope fell below 50%. Because of limited sampling, the exact date at which a patient’s reactive epitope response fell below the 50% threshold was not actually observed. Instead, Barton et al. (2016a) used a non-linear model to estimate the escape time. In order minimize the uncertainty in these estimates of escape time, we excluded data from patients who had already met the criteria before a patient’s first sample or whose escape date estimated by extrapolation after a patient’s final sample. In addition, because we fitted our model to HIV subtype C data, we only used epitopes whose consensus sequence was the same between subtypes B and C. As a result, we were left with only 14 estimates of escape times. Because these estimated escape times are defined at the epitope level, we compared escape time to the mean site sensitivity *G*′ or *G*′. Similarly, we also compared estimated escape times to the mean log conservation for each epitope 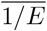.

To test our physico-chemical model’s ability to predict *in vitro* data, we used viral replication rate in cell culture as a measure of HIV fitness. Rihn et al. (2013) estimated the replicative fitness of 31 HIV subtype B viruses bearing mutated capsid CA amino acid residues via spreading replication assay on human MT4 T-cell lines and peripheral blood monocytes. The CA mutants in this study were generated by creating a mutagenized CA library using a low fidelity PCR approach and then inserting the mutated CA sequences in replication competent proviral clones. Fitness was reported as % of the wild-type replication.

### Model Fitting and Inference

The model was fitted to the data using Mathematica 11 (Wolfram Research Inc., 2017). We used the built in, optimization function NMaximize[] with the “NelderMead” option to optimize all of the parameters other than the site specific *G*′. Within each evaluation of NMaximize[], we used FindMaximum[] to identify the optimal 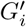 for each site. Using this approach, the model was independently fitted to each region from 200 different initial sets of parameter values. The first 100 fittings used initial values generated by fitting the model to only variable sites and a uniform hierarchical distribution for *G*′. For each of these fittings the estimated parameter values were multiplied by uniform random numbers to further vary their starting values. For the second 100 fittings, the initial values were generated using different random seeds for each run of NMaximize[].

We calculated the Fisher Information Matrix for the 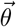 for each region and used it to calculate the 95% Confidence Intervals (CI) for each of its parameter following Bolker (2008, p.197-200). We used these CI to quantify our uncertainty in our estimates and to compare parameters across regions. If the CI for the same parameter in two regions do not overlap, we concluded that the *p* — value for the hypothesis the regions shared the same parameter values was < 0.05.

In order to evaluate how well our physico-chemical model’s site sensitivity *G*′ and the Shannon entropy model’s site conservation 1/*E* predicted available *in vivo* and *in vitro* fitness measures. Neither our *G*′ or 1/*E* estimates nor our empirical *in vivo* and *in vitro* fitness estimates appear normally distributed; therefore, we used Kendall’s non-parametric rank correlation τ to measure the association between our consistent stabilizing selection metrics and the empirical data. Significance tests for our τ estimates were done using Mathematica’s KendallTauTest[]. Test statistics were generated by bootstrapping the data 10,000 times with replacement via the “Permutation” option.

## Results

Briefly, we find that our population genetics and physico-chemical based site sensitivity terms *G*′ are well correlated with the more commonly used entropy based conservation metric 1/*E*. In terms of fitting the data, the entropy model, a saturated parameter model, has an astronomically better AIC score than our best fitting version of our physico-chemical model where Grantham weights and the distribution of site sensitivities vary by region (ΔAIC = 230,007). This poor performance of our model is discouraging, but is not surprising given the fact that the entropy model is a purely descriptive model and fits one parameter for every unconstrained data bin. As a result, the entropy model, by definition, produces the largest *L* value possible given the data. This flexibility comes the cost, in terms of AIC, of a large number of estimated parameters: 19/site × 462 sites = 8778 parameters. This cost, however, is mitigated by the fact that we have more than a 1000 observations for each site. In contrast to its poorer AIC value, we find that our physico-chemical model does a better job predicting empirical *in vivo* and *in vitro* measurements of HIV fitness, if only slightly so. Further, our parameter estimates can be used to test more refined hypotheses such as whether the properties of natural selection, as described by our physico-chemical weights 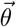 and *G*′ values, varies between protein regions. Thus, while our physico-chemical model appears to capture important biological information embedded in the data, there is clearly much room for future improvement.

### Regional Variation in Model Parameters

The variation and uncertainty in our estimates of 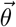 can be found in Table 2 and Figure S1. Although allowing the physico-chemical weights 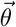 and *G*′ distributions to vary between poly-peptide regions required the addition of just 12 additional parameters and vastly improved the ability of our physico-chemical model to fit the sequence data by 1412 log-likelihood units 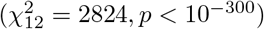. Surprisingly, despite this vast improvement in model fit, the differences in 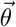 were actually quite small. Further, given the sensitivity of our *L* function to 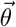, it is perhaps a bit surprising that the choice of a LogNormal or Gamma distribution for *G*′ had no discernible effect on our estimates of a region’s 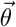. This sensitivity is also reflected by fact that our CI for these parameters are on the order of < 1%. These results clearly indicate that the effects of amino acid substitutions vary between protein regions. Consequently, all of the results we discuss below come from the model fit where 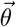 varies between proteins.

**Table 2.**
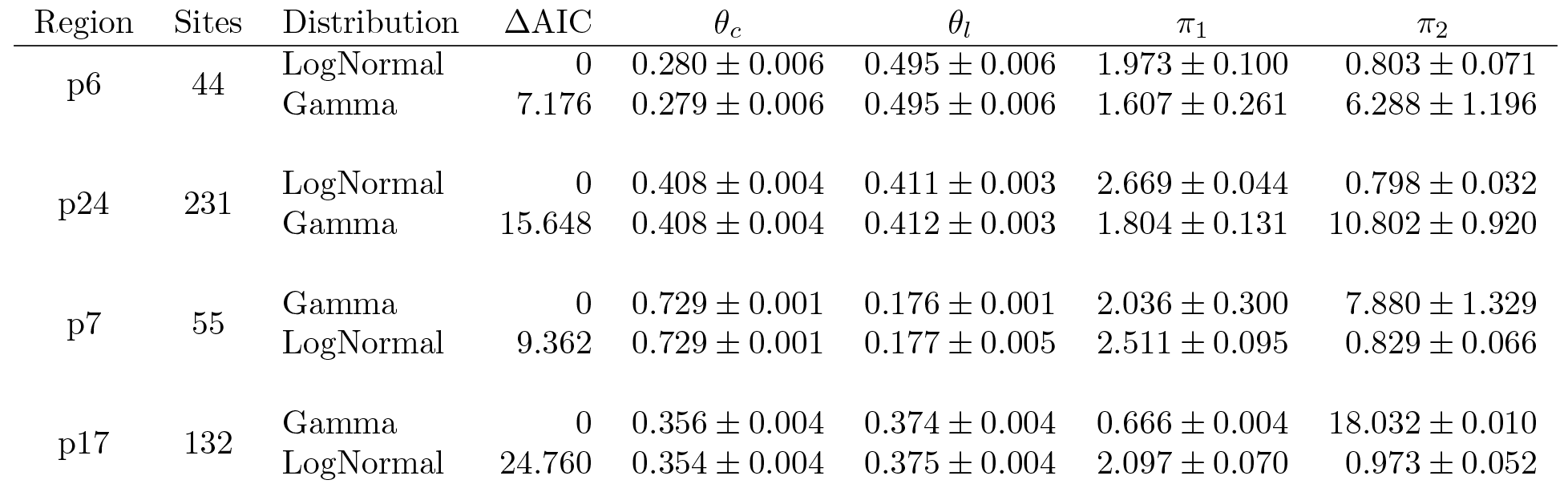
Differences in AIC values AAIC and MLE ±95%CI of physico-chemical model parameters for eac poly-peptide region under Log-Normal and Gamma distributions of site sensitivities *G*′. Note that becaus both distributions have two parameters, their ΔAIC values are simply twice their differences in *L*. The parameters *θ*_*c*_ and *θ*_*l*_, represent physico-chemical model weights for residue composition and polarity, respectively. By definition, *θ*_*v*_ = 1 — (*θ*_*c*_ + *θ*_*l*_) and is not presented. For *G*′ ~ LogNormal(*μ* = π_1_, a = π_2_) an for *G*‣ ~ Gamma(*α* = *π*_1_, *β* = *π*_2_) where *β* is the rate parameter. 95%CI calculated using hessian of LLik surface at MLE (Bolker, 2008, p.197-200).

| Region | Sites | Distribution | $\Delta\text{AIC}$ | $\theta_c$ | $\theta_l$ | $\pi_1$ | $\pi_2$ |
| --- | --- | --- | --- | --- | --- | --- | --- |
| p6 | 44 | LogNormal | 0 | $0.280 \pm 0.006$ | $0.495 \pm 0.006$ | $1.973 \pm 0.100$ | $0.803 \pm 0.071$ |
| | | Gamma | 7.176 | $0.279 \pm 0.006$ | $0.495 \pm 0.006$ | $1.607 \pm 0.261$ | $6.288 \pm 1.196$ |
| p24 | 231 | LogNormal | 0 | $0.408 \pm 0.004$ | $0.411 \pm 0.003$ | $2.669 \pm 0.044$ | $0.798 \pm 0.032$ |
| | | Gamma | 15.648 | $0.408 \pm 0.004$ | $0.412 \pm 0.003$ | $1.804 \pm 0.131$ | $10.802 \pm 0.920$ |
| p7 | 55 | Gamma | 0 | $0.729 \pm 0.001$ | $0.176 \pm 0.001$ | $2.036 \pm 0.300$ | $7.880 \pm 1.329$ |
| | | LogNormal | 9.362 | $0.729 \pm 0.001$ | $0.177 \pm 0.005$ | $2.511 \pm 0.095$ | $0.829 \pm 0.066$ |
| p17 | 132 | Gamma | 0 | $0.356 \pm 0.004$ | $0.374 \pm 0.004$ | $0.666 \pm 0.004$ | $18.032 \pm 0.010$ |
| | | LogNormal | 24.760 | $0.354 \pm 0.004$ | $0.375 \pm 0.004$ | $2.097 \pm 0.070$ | $0.973 \pm 0.052$ |

In terms of describing the distribution of site sensitivities *G*′ across a given poly-peptide region, the LogNormal distribution performed substantially better than the Gamma distribution with the p6 nucleo protein and p24 matrix protein (ΔAIC = 7.2 and 15.6, respectively). The LogNormal distribution mean parameter *μ* was indistinguishable between these two regions, but the standard deviation parameter σ was significantly lower in p6 than p24. In contrast, the Gamma distribution performed substantially better than the LogNormal distribution for the p7 nucleocapsid protein and p17 capsid protein (ΔAIC = 9.4 and 24.8, respectively). The Gamma distribution shape parameter *α* was indistinguishable between these two regions, but the rate parameter 0 was significantly lower in p7 than p17. Taken together, these results indicate that the distribution of *G*′ values varies between protein regions. This is in spite of the fact that the first two central moments of p6, p7, and p17 are statistically indistinguishable (Fig. 1 and Table S1).

**Fig 1.**
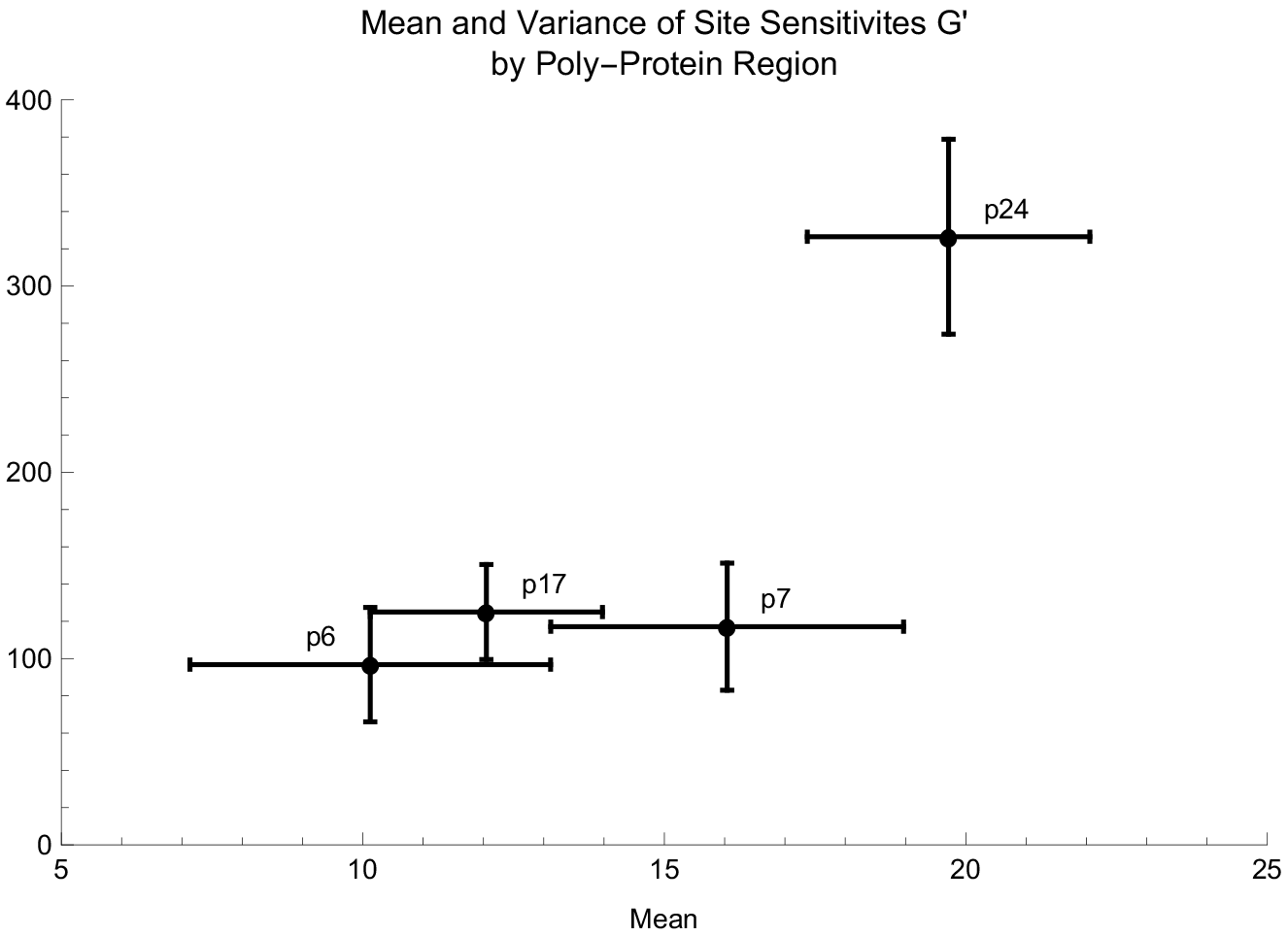
Comparison of estimates of mean and variance of *G*′ for poly-peptide regions. Error bars represent parameter 95% CI. CI estimated using the t distribution and the likelihood ratio test squares after Zar (1999, p. 99 and 111). Note that even thought the central moments of p6, p7, and p17 are statistically indistinguishable, the model parameters describing their distribution are distinguishable.

### Site Sensitivity *G*′ vs. Site Conservation 1/*E*

Kendall’s rank correlation τ between our model’s site sensitivity parameter *G*′ and the entropy model’s site conservation 1/*E* metric indicate that they are well correlated with one another (τ = 0.598 — 0.720 with *p* < 10^−10^ for all regions, Fig 2).

**Fig 2.**
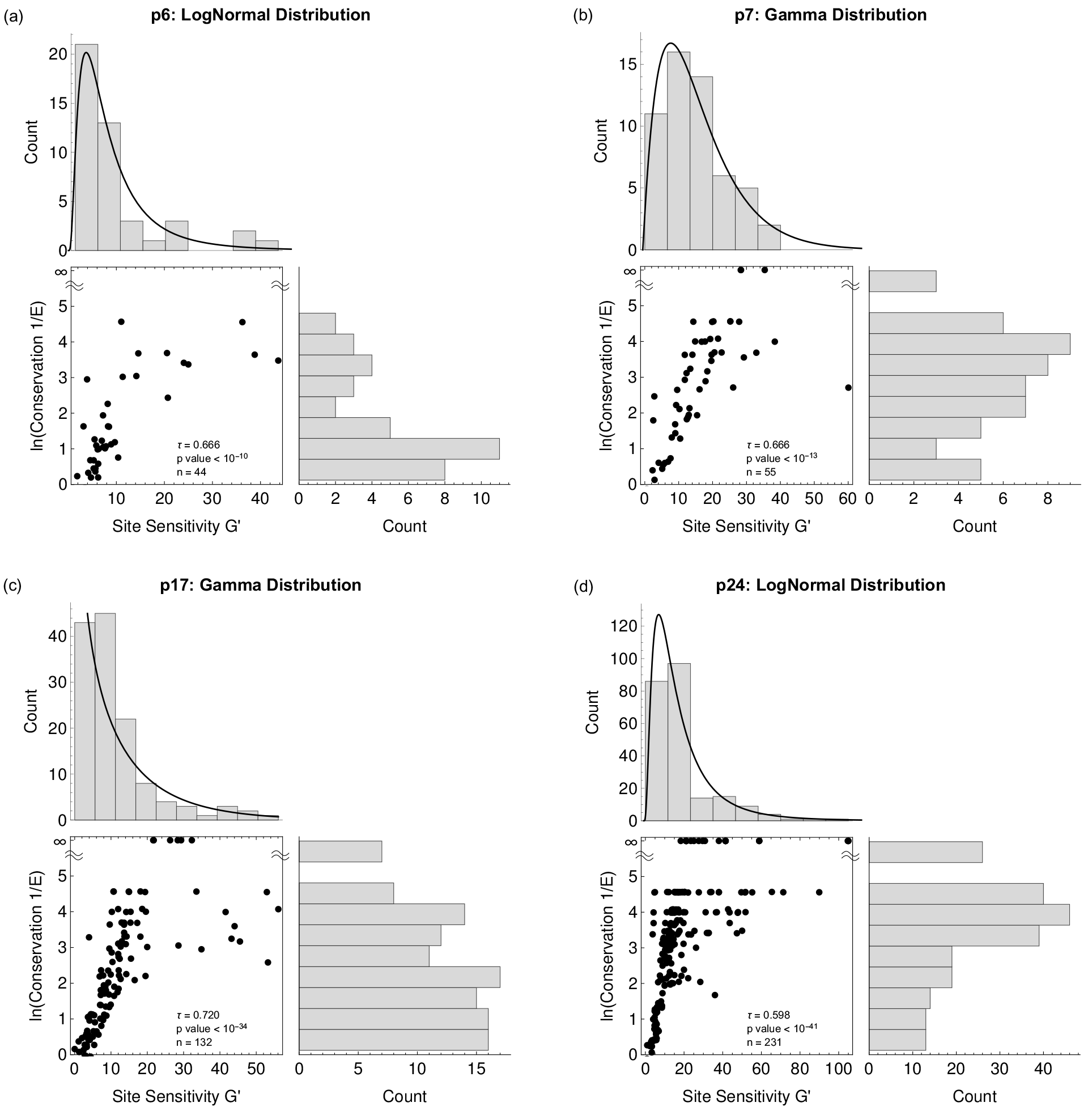
Comparison of site specific physico-chemical sensitivity *G*′ and site conservation 1/*E* for each Gag poly-peptide region using Kendall τ correlation statistic. Invariant sites, by definition, have a 1/*E* value of ∞. In contrast, because our model assumes *G*′ is pulled from either a LogNormal or Gamma probability distribution whose parameters are estimated simultaneously with *G*′, invariant sites have finite *G*′ values. *G*′ values, their distribution, and the parameters used are the best fitting model for each region (Table 2). Regions: (a) nucleo protein p6 with *G*′ ~ Gamma, (b) nucleocapsid protein p7 *G*′ ~ LogNormal, (c) matrix protein p17 *G*′ ~ LogNormal, and (d) capsid protein p24 *G*′ ~ Gamma. Note, *n* is the number of sites fitted for each region.

### Predicting Empirical HIV Fitness Data

As expected, both site conservation 1/*E* and site sensitivity *G*′ were positively correlated with the *in vivo* measure of fitness: escape time (τ = 0.291 and 0.475, respectively; Fig 3 and Table S2). However, only the correlation between *G*′ and escape time was significant (*p* = 0.016 vs. 0.141 for 1/*E*).

**Fig 3.**
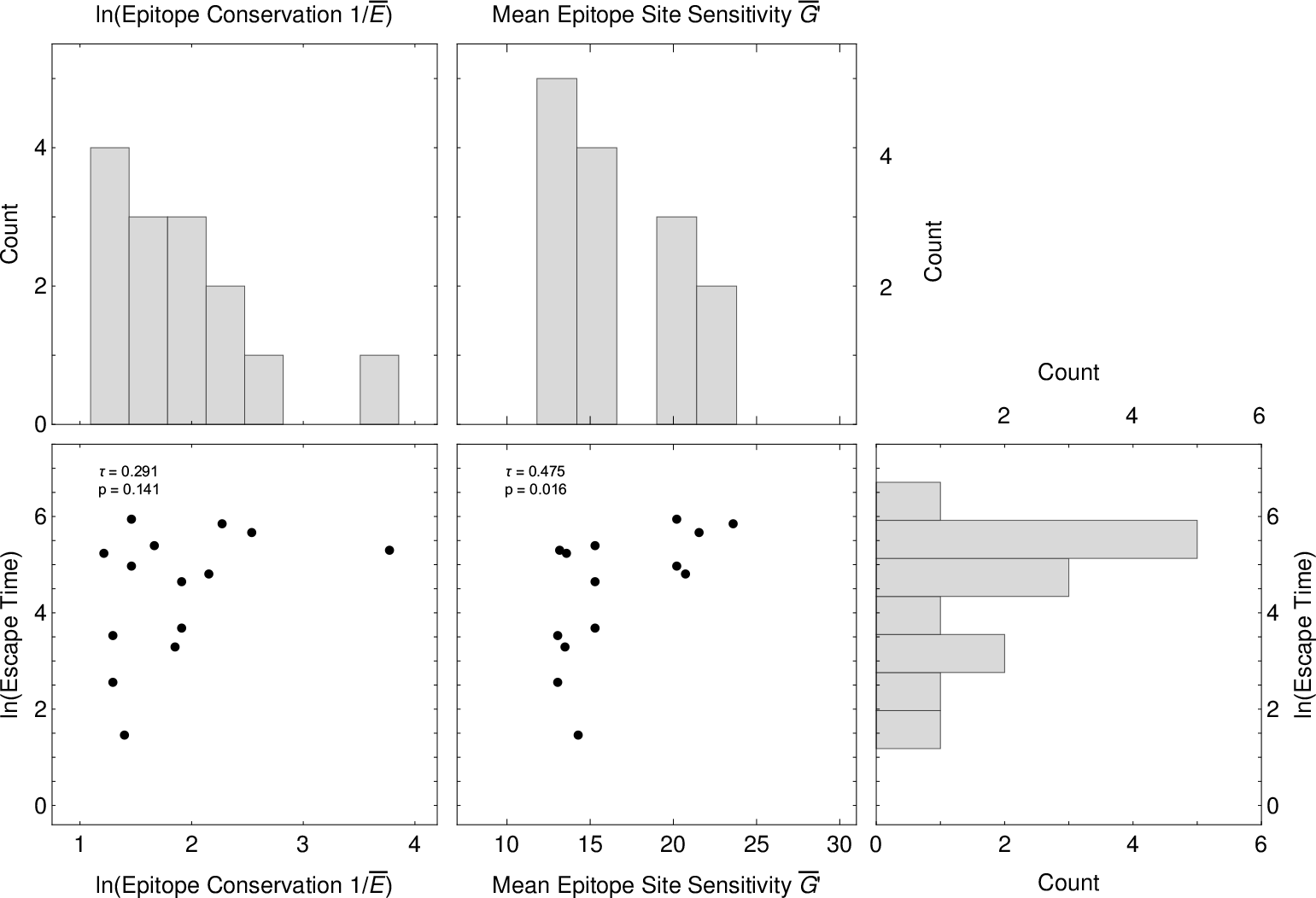
Kendall rank correlation τ of mean log conservation ln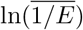 and mean site sensitivity 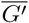 with ln(escape time) of 14 epitopes of the Gag poly-peptide whose immune escape was observed (original data from Liu et al. (2013) results from Barton et al. (2016b) used here). Correlation *p* values estimated by bootstrapping data 10,000 times.

Both *G*′ and 1/*E* were also positively correlated with changes to the capsid protein on *in vitro* replication fitness (τ = 0.238 and 0.211, respectively; Fig. 4). However, as with the escape time predictions, only the *G*′’s t was significant (*p* = 0.047 vs. 0.080 for 1/*E*). Similarly, *G*′ and 1/*E* were also positively correlated with changes to the capsid protein on *in vitro* spreading fitness (τ = 0.231 and 0.157), but neither predictor’s τ was significantly greater than 0 (*p* = 0.055 and 0.192, respectively).

**Fig 4.**
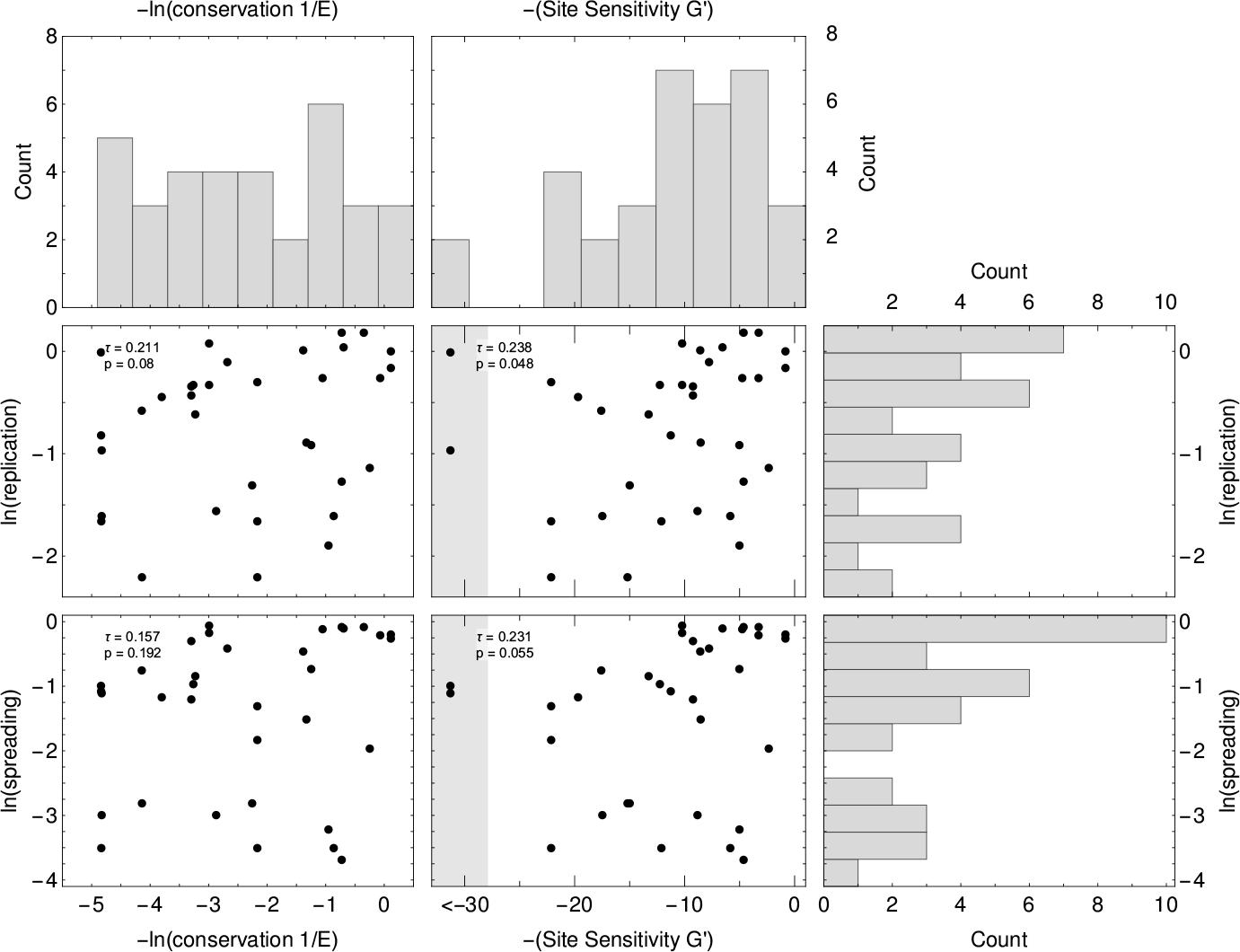
Metrics vs Spreading fitness. Correlation of conservation and sensitivity of residues with spreading fitness of 31 viral strains bearing mutations in the corresponding residues A) Correlation of the assayed spreading fitness of the mutated virus in days with the correspondingly position’s entropy B) Correlation of the assayed spreading fitness of the mutated virus in days with the corresponding position’s physico-chemical sensitivity.

## Discussion

Although initially introduced as a metaphor for describing how evolution works, fitness landscapes have proven to be useful tools in theoretical and empirical research (for example, Kauffman, 1993; Wilke and Drummond, 2006; Gilchrist, 2007; Gilchrist et al., 2009; Wallace et al., 2013; Gilchrist et al., 2015, and citations below). Ideally, HIV fitness landscapes can help researchers predict when and where a virus will escape immune control (Barton et al., 2016a) and develop effective vaccines (Ferguson et al., 2013; Shekhar et al., 2013). In a few cases HIV fitness landscapes have been estimated directly from experimental data (Hinkley et al., 2011; Kouyos et al., 2012; Mann et al., 2014; Rihn et al., 2013), but in most studies the landscape is inferred from the ever growing libraries of genotype frequency data (Barton et al., 2016a; Zanini et al., 2016; Mann et al., 2014; Ferguson et al., 2013; Deforche et al., 2008; Seifert et al., 2015).

In this study we utilize fundamental findings from the field of population genetics to infer HIV Gag’s poly-peptide physico-chemical fitness landscape using count data from the LANL HIV database. In contrast to explicitly mapping out HIV’s fitness landscape, other researchers have used site conservation 1/*E*, where *E* is the Shannon entropy of a site, as a proxy for the strength of consistent stabilizing selection (Rihn et al., 2013; Ferrari et al., 2011; Liu et al., 2012b). While this interpretation may seem intuitive, 1/*E* is actually a summary statistic rather than a measure of the selection coefficient between an optimal amino acid and its alternatives. Further, entropy based metrics such as 1/*E* result from fitting saturated models, where the number of parameters is on par with the number of data categories. As a result, it is perhaps not surprising that the AIC for the entropy model fit is still substantially better than its value for our highly unsaturated physico-chemical model (ΔAIC = 230,007). We note that because, 1/*E* = ∞ at invariant sites, 1/*E* for an invariant site will change dramatically if new data includes even a single alternative amino acid at that site. In contrast, because we assume *G*′ comes from a probability distribution whose parameters are estimated simultaneously with our *G*′ values, the metric will not change as dramatically if an alternative amino acid is eventually observed at that site.

While saturated models have the advantage of being maximally flexible in terms of fitting data, this flexibility comes at the cost of being minimally informative about the processes generating the data. For example, our physico-chemical model’s site sensitivity *G*′ did a better job than the entropy model’s site conservation 1/*E* in predicting *in vivo* and *in vitro* HIV fitness measurements (Figs 3 and 4). These results suggest that our physico-chemical model is more efficient at extracting biological meaningful information from the sequence data we used to fit our models.

In addition to extracting more meaning from the data, because our physico-chemical model is derived from biological principles, it can be used to evaluate biologically based hypotheses. For example, we find that the impact of our three different physico-chemical traits 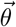 on fitness varies substantially between between protein regions (Table 2). These results indicates the effects of substituting one amino acid residue for another varies with its broader genetic context. This is, perhaps, unsurprising given that previous researchers have found that interior regions of the protein are more sensitive to changes in polarity (Nakai et al., 1988). Nevertheless, our ability to detect these slight regional differences in the character of the consistent stabilizing selection illustrates the statistical power of our physico-chemical modeling approach.

In addition, site sensitivities *G*′ values differ between protein regions in terms of the distribution that best describe the estimated values (LogNormal vs. Gamma). The fact that the *G*′ values of different protein regions follow different distributions suggests that the contribution of each site to protein function varies between sites. For example, the LogNormal distribution suggests the *G*′ values are the result many positive random independent variables acting in a multiplicative manner. In contrast, the Gamma distribution suggests that protein function depends on fluxing through different conformational states with exponential like waiting times for each state. Even amongst the p6 and p24 regions where site sensitivities *G*′ are best described by a LogNormal distribution, they appear to come from distributions with different *μ* parameters (Table 2). For example, the average *G*′ values for region p24 were greater than p7 and p17, which is consistent with previous findings (Rihn et al., 2013; Martinez-Picado et al., 2006) (Figure 1). Despite our ability to differentiate between different hypothesized distributions for *G*′, our estimates of physico-chemical weightings 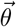 are actually insensitive to these distributional differences (Table 2). One plausible hypothesis for these peptide region specific differences in *G*′ distributions could be their differences in secondary structure. This idea could be tested by determining whether grouping sites by secondary structure (or some other feature such as distance from surface or active site) provides a better fit than grouping sites by protein region as we do here.

While our physico-chemical model improves our ability to predict empirical HIV fitness data, there is still a substantial amount of noise in our predictions. This variation likely has a number of different sources. In terms of the data we are trying to predict, the *in vitro* spreading fitness measures suffer from the unnatural qualities of cell culture. Similarly, the *in vivo* epitope escape fitness measures likely includes substantial effects due to biological variation between patients, is based on a limited number of sample time points, and limited sequencing depth to determine genotype frequencies.

In terms of model shortcomings, there are many. For example, our choice of physico-chemical properties to include in our distance function was based solely on Grantham’s classic work (Grantham, 1974). Fortunately, testing the power of other physico-chemical properties is straightforward with our model. Further, in its current form, our physico-chemical model ignores the effect of mutation bias. Mutation bias, i.e. the fact that mutation rates between residues differ from one another, can also contribute to the probability of observing a particular codon and, in turn, amino acid. Although the effects of mutation bias are likely to be unimportant at sites with large site sensitivities *G*′, mutation bias can dominate the evolutionary outcome of sites under weak selection.

Moving on to shortcomings that are more challenging to over come, both the physico-chemical and entropy models assume statistical independence between patient samples. These assumptions also shared by the more complex Ising and Potts models and could be addressed by extending our approach to a phylogenetic framework (Beaulieu et al., In Review). Both the physico-chemical and entropy models also assume a single, invariant fitness peak centered around a site’s optimal amino acid residue *a*_*_. While it should be possible to extend our physico-chemical to allow for more than one peak in the amino acid residue landscape by modifying our distance function, we suspect our ability to reject the simpler hypothesis of a single peak would be weak, especially if they occurred in similar points in physico-chemical space. Allowing *a*_*_ to explicitly switch over time, would be even more challenging than allowing for multiple fitness peaks. We expect detecting such switches would be very difficult without large, high quality and high resolution datasets.

Perhaps most glaringly shortcoming is the fact that our physico-chemical ignores epistatic effects. Epistatic interactions are likely ubiquitous and have been well documented in the virus literature (Koek et al., 2012; Brockman et al., 2007, 2010). Fortunately, we could extend our physico-chemical model to include the effects of epistatic interactions between sites in a similar manner to the Ising and Potts model. In the same way that we use a physico-chemical distance function and site sensitivity *G*′ to generate the predicted frequency of the 20 canonical amino acid residues, we could define a more complex distance function that would allow us to describe epistatic effects between sites and predict the probability of the 400 different possible site pairs of amino acid residues using a relatively small number of parameters. The end result would ideally be a more realistic model than the Ising model, but one that requires many fewer parameters, is more biologically informative, and is easier to fit than the Potts model.

In conclusion, we argue that one promising way to improve our ability to extract biologically meaningful information from sequence databases is to use well defined and biologically grounded models. Here we show how our physico-chemical model can improve our ability predict HIV fitness. Because our model is grounded in the field of population genetics, it is inherently well suited to describe evolutionary data. Because our model is well defined biologically, it provides a clear framework to test specific hypotheses about HIV protein sequence evolution and can serve as the basis for more complex, but parameter limited, models.

## Acknowledgements

M.A.G received financial support from NSF grants MCB-1546402 (Primary Investigator: A. VonArnim), MCB-1120370 (Primary Investigator: M.A.G), and DEB-1355033 (Primary Investigator: Brian O’Meara) as well as the Department of Ecology and Evolutionary Biology at the University of Tennessee and the National Institute for Mathematical and Biological Synthesis (NIMBioS, NSF:DBI-1300426 with additional support from the University of Tennessee). E.G.H. received financial support from the Departments of Mathematics and Microbiology at the University of Tennessee and fellowship from NIMBioS. We also gratefully acknowledge assistance from Dr. Vitaly V. Ganusov.

